# Spike mutation resilient scFv76 antibody counteracts SARS-CoV-2 lung damage upon aerosol delivery

**DOI:** 10.1101/2022.05.27.493569

**Authors:** Ferdinando M. Milazzo, Antonio Chaves-Sanjuan, Olga Minenkova, Daniela Santapaola, Anna M. Anastasi, Gianfranco Battistuzzi, Caterina Chiapparino, Antonio Rosi, Emilio Merlo Pich, Claudio Albertoni, Emanuele Marra, Laura Luberto, Cécile Viollet, Luigi G. Spagnoli, Anna Riccio, Antonio Rossi, M. Gabriella Santoro, Federico Ballabio, Cristina Paissoni, Carlo Camilloni, Martino Bolognesi, Rita De Santis

## Abstract

Uneven worldwide vaccination coverage against SARS-CoV-2 and emergence of variants escaping immunity call for broadly-effective and easily-deployable therapeutics. We previously described the human single-chain scFv76 antibody, which recognizes SARS-CoV-2 Alfa, Beta, Gamma and Delta variants. We now show that scFv76 also neutralizes infectivity and fusogenic activity of Omicron BA.1 and BA.2 variants. Cryo-EM analysis reveals that scFv76 binds to a well-conserved SARS-CoV-2 spike epitope, providing the structural basis for its broad-spectrum activity. Moreover, we demonstrate that nebulized scFv76 exhibits therapeutic efficacy in a severe hACE2 transgenic mouse model of COVID-19 pneumonia, as shown by body weight and pulmonary viral load data. Counteraction of infection correlates with the inhibition of lung inflammation observed by histopathology and expression of inflammatory cytokines and chemokines. Biomarkers of pulmonary endothelial damage were also significantly reduced in scFv76-treated mice. Altogether the results support the use of nebulized scFv76 for COVID-19 induced by any SARS-CoV-2 variants emerged so far.

## INTRODUCTION

Lung infection from emerging viruses can raise serious public health concern in case of pandemics. From the last coronavirus disease 2019 (COVID-19) pandemic, caused by the severe acute respiratory syndrome coronavirus 2 (SARS-CoV-2), we learned how a broad and timely vaccination campaign, together with the adoption of prevention measures like mask wearing and social distancing, and with the use of antiviral medications, can reduce deaths and intensive care pressure. The relatively milder disease, recently associated to the emergence of the Omicron BA.1 and BA.2 variants, is raising hopes for a weakening of the pandemic (*1*). However, because of the uneven worldwide vaccination coverage and possible emergence of new viral variants escaping immunity, the evolution of COVID-19 is unpredictable and re-occurrence of severe pulmonary diseases cannot be ruled out (*2*). Moreover, the observation of several threatening post-acute sequelae of SARS-CoV-2 infection affecting particularly the nervous and cardiovascular systems (*3*), is urgently calling for easily deployable therapeutic measures able to timely control the infection in the early stages. In a prospective COVID-19 pandemic re-exacerbation, and even in the case of transition into an endemic phase, two types of intervention are being envisaged: first, improve vaccine equity worldwide with a possible update against SARS-CoV-2 variants and second, validate early-stage therapeutic protocols preventing worsening of the disease and ultimately hospitalizations and post-acute sequelae. As of today, Omicron variants are challenging the efficacy of most injected antibodies (*4-10*). Moreover, because the Omicron variants apparently remain confined mainly to the upper respiratory tract (*11*), the use of systemic antibodies is becoming somehow questionable.

We recently described a cluster of human anti-SARS-CoV-2 antibodies in the format of single-chain variable fragment (scFv) able to neutralize viral variants both *in vitro* and in animal models (*12*). We also showed that such antibody format is suitable for intra-nasal or aerosol formulations that might be useful for the topical treatment of upper and lower respiratory tract SARS-CoV-2 infections (*12*).

In the present work, we show that the scFv76 antibody of the cluster, previously found able to react with SARS-CoV-2 Alfa, Beta, Gamma and Delta, is also resilient to the Omicron BA.1 and BA.2 mutations substantially retaining neutralizing activity against these new viral variants. Moreover, we provide pre-clinical proof of concept of the nebulized scFv76 efficacy in a mouse model of Delta infection, selected as an aggressive prototype of viral pneumonia. Finally, we prove, by single particle Cryo-EM, the wide recognition properties of the scFv76 antibody at the molecular level showing that it binds to a well conserved epitope at the tip of the spike protein within the receptor binding domain (RBD), with an architecture that is able to accommodate the mutations found in all SARS-CoV-2 variants known to date. Our results support the use of scFv76 antibody for aerosol therapy of COVID-19 induced by all variants of concern.

## RESULTS

### ScFv76 efficiently neutralizes SARS-CoV-2 Delta and Omicron

The scFv76 antibody was previously described to be able to neutralize the SARS-CoV-2 Alpha, Beta, Gamma and Delta viral variants both *in vitro* and in animal models (*12*). To evaluate its reactivity with the recently emerged Omicron variants, the ability to compete the binding of the Omicron BA.1 and BA.2 spikes to human ACE2 was tested by ELISA. Results in **Table 1** show that scFv76 can inhibit Omicron BA.1 and BA.2 spike binding to ACE2 at IC50 concentrations <2.5 nM, which is similar to the potency against Delta. Binding affinity of scFv76 to the Omicron spikes was then tested by Surface Plasmon Resonance (SPR) showing K_D_ of 6.3 and 14.5 nM for BA.1 and BA.2, respectively (**Table 2**). Neutralizing activity against SARS-CoV-2-S Omicron BA.1 and BA.2 pseudotyped viruses was also exhibited by scFv76, but not by scFv5 (an anti-RBD antibody previously shown to be devoid of neutralizing activity and used as negative control) (*12*), with an IC50 value of 2.84 and 2.47 nM, respectively (**Figure 1A**).

**Table 1.** ScFv76 spike/ACE2 competition by ELISA

| | IC50 nM (±SE) | | |
| --- | --- | --- | --- |
|  | Delta | Omicron BA.1 | Omicron BA.2 |
| scFv76 | 1.64 (0.25) | 1.90 (0.3) | 2.4 (0.1) |
| scFv5 | >40 | >40 | >40 |

Competition of spike S1 binding to human ACE2 by scFv antibodies measured by ELISA. IC50 values (expressed as nM concentration) are the average ( ± SE) from 3-4 independent experiments.

**Table 2.** ScFv76 Surface Plasmon Resonance (SPR) data

| | Spike Trimer | $k_a$<br>($10^5 \text{ M}^{-1} \text{ s}^{-1}$) | $k_d$<br>($10^{-5} \text{ s}^{-1}$) | $K_D$<br>(nM) |
| --- | --- | --- | --- | --- |
| scFv76 | Omicron BA.1 | 0.7 | 41.6 | 6.3 |
|  | Omicron BA.2 | 1.2 | 174.9 | 14.5 |

**Figure 1.**
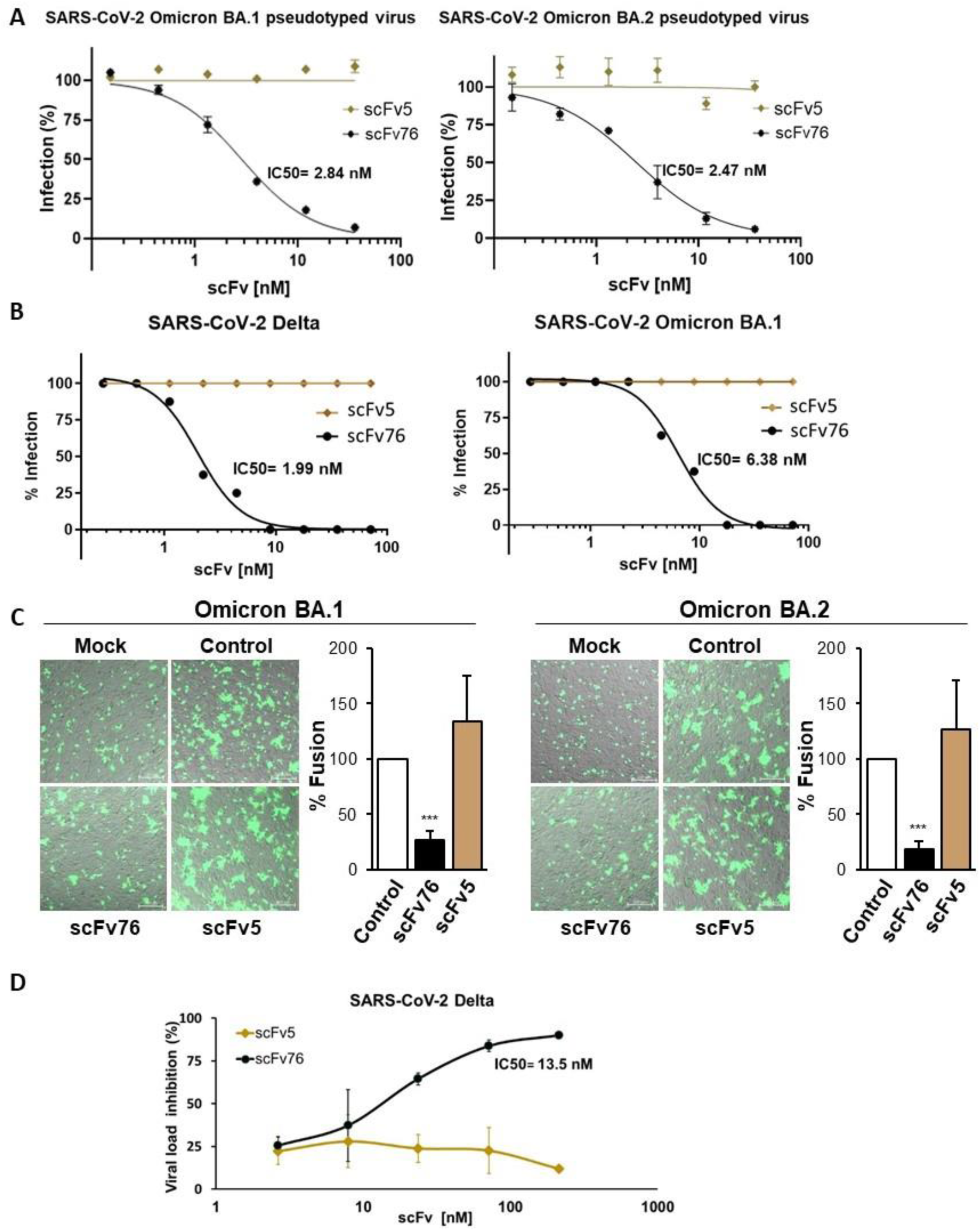
Resilience of scFv76 reactivity to Omicron BA.1 and BA.2 variants. A) Neutralization of pseudotyped-virus expressing the SARS-CoV-2 Omicron (B.1.1.529) BA.1 or BA.2 spike assessed by luciferase-assay in hACE2-expressing Caco-2 cells. Data are the average (±SD) of two replicates from one representative experiment. B) Neutralization activity of scFv antibodies assessed by viral titration (Delta and Omicron strains) on Vero E6 cells by microneutralization assay. Data are the average (±SD) of eight replicates from one representative experiment. C) Inhibition of SARS-CoV-2 spike-mediated cell-cell fusion using HEK293T donor cells expressing GFP and Omicron BA.1 or BA.2 spike, or GFP only (mock), incubated 1 h with scFv76 or scFv5 (360 nM) and then overlaid on monolayers of hACE2-expressing A549 cells for 24 h. The overlay of bright-field and fluorescence images is shown. Scale bar, 200 nm. Cell-cell fusion quantification is expressed as percentage relative to control (average ± SD of 5 fields from two biological replicates). ***p < 0.001 (ANOVA). D) Neutralization of the authentic SARS-CoV-2 Delta virus in Calu-3 cells. Serially diluted (3-fold) antibodies were added to cells 1 h after infection. Quantification of viral load was done by RT-qPCR 72 h after infection. Data are the average (±SD) of two independent experiments. The IC50 value (expressed as nM concentration) is also shown in A), B) and D) panels.

Neutralization of infectivity was further tested against authentic SARS-CoV-2 Delta and Omicron BA.1 virus by microneutralization assay of cytopathic effect (CPE) in Vero E6 cells. In this assay, scFv76 exhibited IC50 values of 1.99 and 6.38 nM against the Delta and Omicron BA.1 variant, respectively, whereas the not-neutralizing antibody scFv5 showed no anti-viral activity, as expected (**Figure 1B**).

The Omicron BA.2 spike was recently shown to be more pathogenic and more efficient in mediating syncytia formation than the BA.1 spike (*13*). The ability of scFv76 to prevent SARS-CoV-2 Omicron BA.1 or BA.2 spike-induced fusion of pulmonary cells was than tested *in vitro*. As shown in **Figure 1C**, incubation with nanomolar concentrations of the scFv76 antibody proved to be significantly effective at inhibiting fusion between both BA.1 and BA.2 spike-expressing human HEK293T cells and human lung A549 cells stably expressing the hACE2 receptor (A549 hACE2).

Preliminarily to an *in vivo* pharmacology study of nebulized scFv76 in a severe Delta-induced pneumonia mouse model, its antiviral neutralization potency was tested *in vitro* by RT-qPCR in Delta-infected pulmonary Calu-3 cells in comparison with the not-neutralizing control antibody scFv5. As shown in **Figure 1D**, scFv76 was found to inhibit infection with an IC50 of 13.5 nM, whereas no activity of the control antibody at concentration >200 nM was observed.

### Therapeutic efficacy of nebulized scFv76 in a severe SARS-CoV-2 Delta interstitial pneumonia model

We previously established the biochemical suitability of scFv76 to aerosol delivery by a mesh nebulizer (*12*). To test pharmacological efficacy of the aerosolized antibody, pneumonia infection was established in transgenic hACE2 mice by intranasal challenge with 1×10^5^ TCID50 SARS-CoV-2 (Strain Delta B.1.617.2). The overall experimental design is shown in **Figure 2A**. Differently from infected mice treated with vehicle, the group of mice treated with scFv76 showed significant body weight recovery, 4 days post infection (**Figure 2B**). This result correlated with about 100-fold reduction in the lung viral RNA copy number, as assessed by RT-qPCR (**Figure 2C**), and with reduction of infectious viral particles, as measured by TCID50 (**Figure 2D**). Notably, the nebulized scFv76 reduced infectious virus titer in the lungs to undetectable levels in three out of five mice; significant viral RNA reduction was also observed in the nasal turbinates (**Figure 2E**). Consistently, histopathological analysis of lung sections showed significant reduction of lung interstitial edema and hematic endoalveolar extravasation, reduction of cellular inflammatory infiltrates in the alveolar/interstitial space and reduction of alveolar septal thickening (**Figure 3A**). Overall, the treatment with nebulized scFv76, but not PBS, was significantly effective at counteracting the lung inflammation and damage induced by the Delta virus as scored in **Figure 3B**. In order to further evaluate the extent of protection conferred by scFv76 nebulization in Delta infected mice, RT-qPCR analyses were performed to measure the mRNA expression of several inflammatory effectors in lung homogenates, harvested 4 days post infection. Data indicate that the aerosol treatment with scFv76 induced significant reduction of key pro-inflammatory cytokines like IL-6, IL-1β, IL-21, IL-10, IL-4 and TNF-α and chemokines Ccl2, Ccl20, Cxcl-1 and Cxcl-10 (**Figure 4A** and **B**). Notably, lung of infected and vehicle-treated mice showed upregulated transcription levels of both Type-I (IFN-α and, especially, IFN-β) and Type-II (IFN-γ) interferons, as well as that of key IFN-modulated genes (Ifit1, ISG15, MX1). All these genes resulted significantly reduced in lungs of mice treated with scFv76 (**Figures 4A** and **C**). Finally, we evaluated some biomarkers of pulmonary vascular damage and data indicated that the treatment also counteracted upregulation of infection-induced tissue damage molecules including adhesion molecules, angiopoietin 2 and inflammasome effectors such as NLRP3 (**Figures 4D** and **E**).

**Figure 2.**
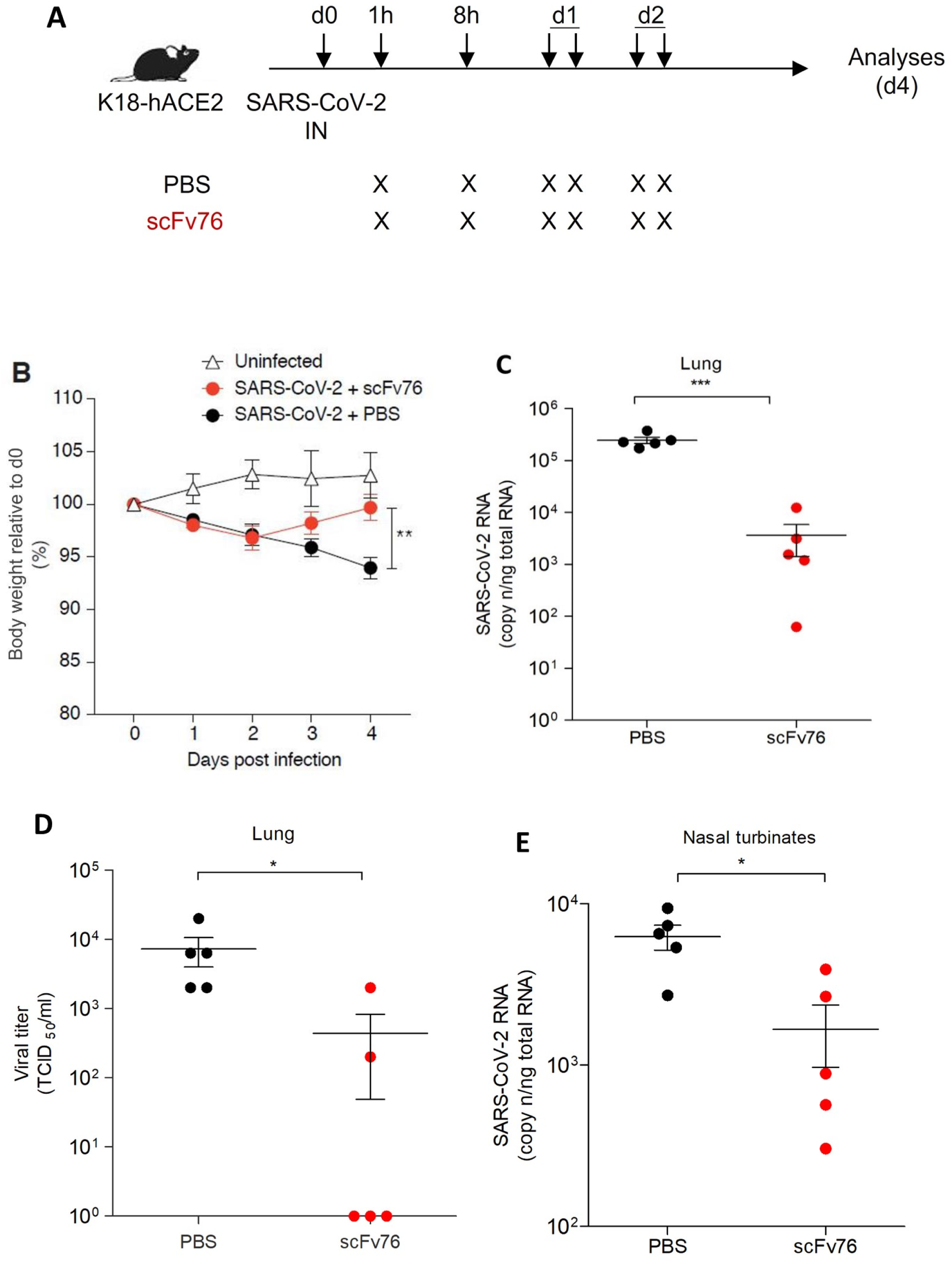
Therapeutic efficacy of nebulized scFv76 in a mouse SARS-CoV-2 Delta pneumonia model. A) Study design. Human ACE2 transgenic mice were nose-only exposed to 2.5 mL of 3 mg/mL scFv76 solution, or PBS (as vehicle-control), 1 h and 8 h after SARS-CoV-2 Delta intranasal infection (1×10^5^ TCID50/mouse), and twice/day for two additional days. B) Body weight changes. Daily body weights from day 0 to day 4 were recorded for each group and plotted as percentage with respect to day 0. Data are the average (±SE). Day on which there was a significant difference in average % body weight between scFv76-treated or PBS-treated animals is denoted by **(p-value < 0.01). C) Lung viral RNA quantification. At day 4 post infection, lungs were collected for viral RNA quantification by RT-qPCR. Each dot represents one mouse. Data are expressed as the copy number/ng total RNA (n=5). D) Lung virus titration. Viral titers in the lung 4 days post infection were determined by viral median tissue culture infectious dose (TCID50) assay. Each dot represents one mouse. Data are expressed as TCID50/mL. E) Viral RNA quantification in nasal turbinates. Quantification by RT-qPCR in nasal turbinates and data representation were as in C). Statistical differences in B), C), D) and E) were assessed by the Mann-Whitney U-test. Significance is indicated as follows: *p<0.05; **p<0.01; ***p<0.001

**Figure 3.**
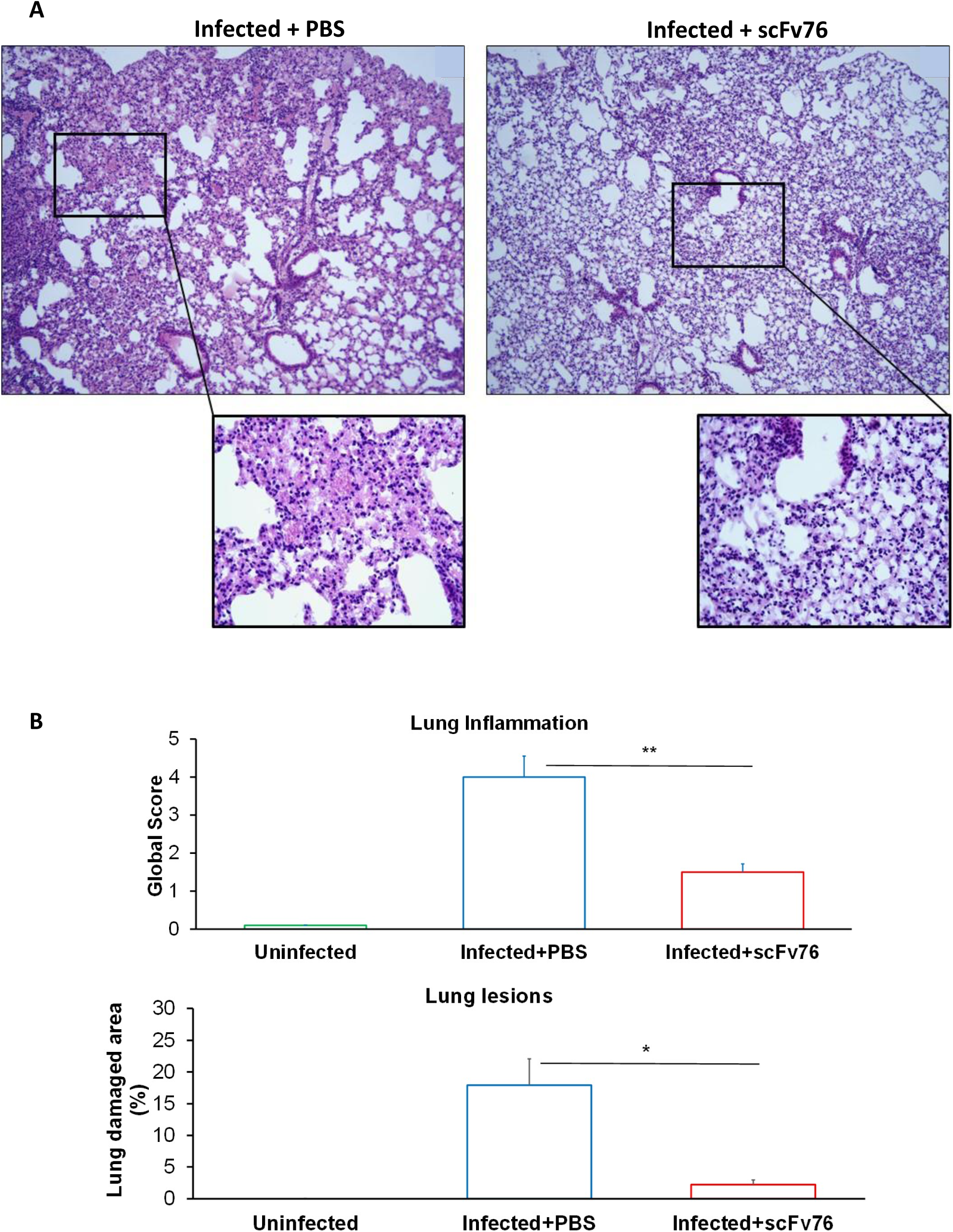
Therapeutic efficacy of nebulized scFv76 correlates with reduction of inflammatory scores. A) Histopathological analysis of lung tissue sections from mice challenged with SARS- CoV-2 Delta and treated by aerosol with scFv76, or PBS, as described in Figure 2. Representative pictures of lung sections, stained with hematoxylin and eosin, from PBS- (left panel) or scFv76-treated (right panel) mice. Overall, scFv76 treatment counteracted the infection-induced diffuse lung edemas and hematic endoalveolar extravasation associated to cellular inflammatory infiltrates in the alveolar/interstitial space and alveolar septal thickening, observed in PBS-treated mice. Scale bar, 200 μm; 10 × magnification. Inset: zoom, 40 × magnification. B) Scores of overall lung inflammation (upper panel) and lung lesion (bottom panel) measured on lung sections as in A). Data are the average (±SE) (n=5) and are expressed as global score and lung damaged area (%), respectively (scoring details in Materials and Methods). Statistical analysis by the Student’s t test. Significance is indicated as follows: * p< 0.05; **p<0.01.

**Figure 4.**
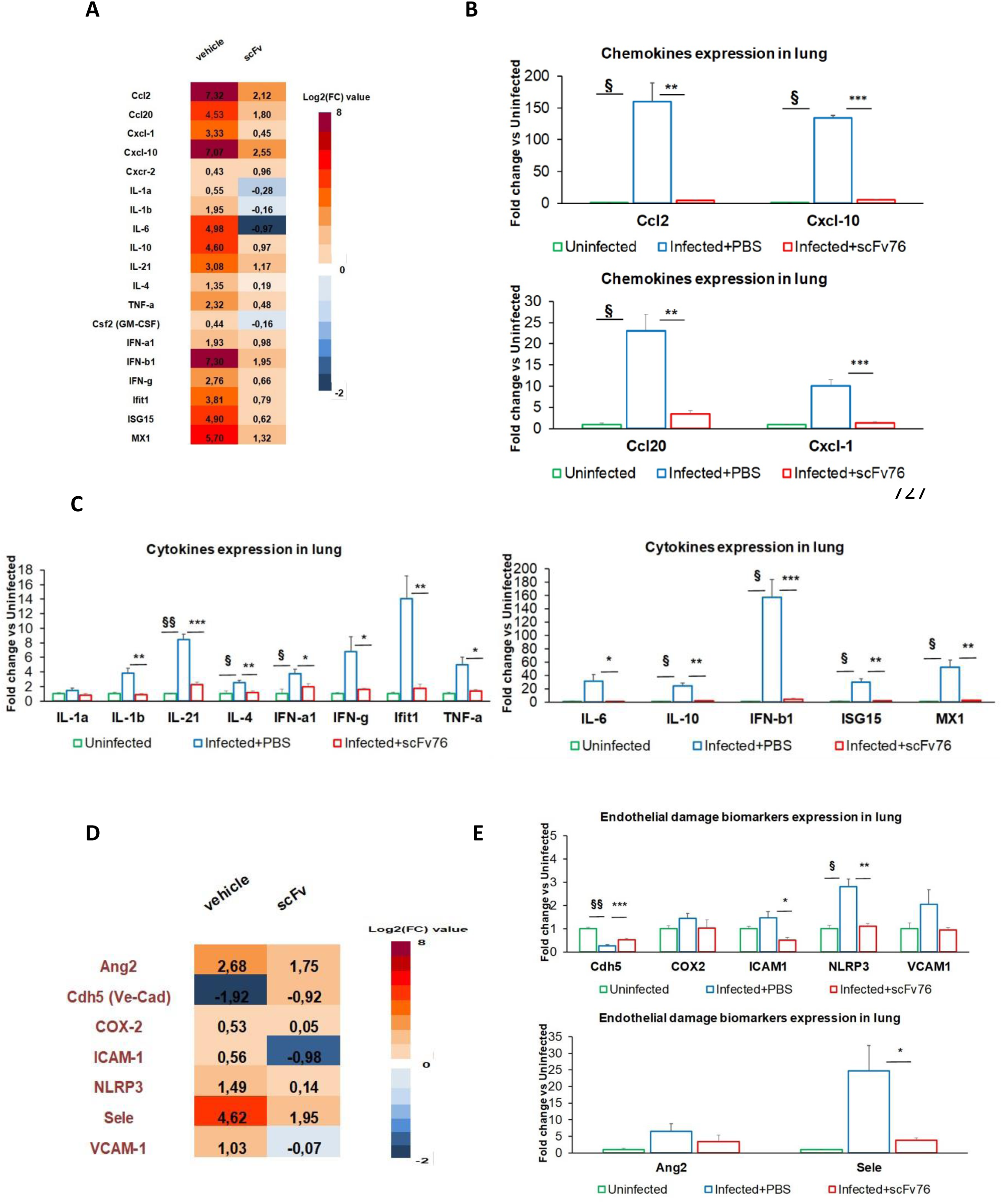
Therapeutic efficacy of nebulized scFv76 correlates with reduction of pulmonary inflammatory and vascular damage biomarkers. A) Heat map of differential gene expression analysis for inflammatory effectors, as determined by Real Time RT-qPCR, in lung homogenates of mice challenged with SARS-CoV-2 Delta and treated by aerosol with scFv76 or PBS. Data are the average of the Log2 expression fold change (FC) obtained from each experimental group (n=5) with respect to uninfected mice. Up-regulation appears as shades of red and down-regulation appears as shades of blue. B) The mRNA expression level of genes encoding for key chemokines assessed by Real Time RT-qPCR in samples as in A). Results are the average (±SE) of expression fold change (FC) with respect to uninfected mice. C) The mRNA expression levels of genes encoding for key inflammatory effectors assessed by Real Time RT-qPCR as above. Results are expressed as in C). D) Heat map of differential gene expression analysis for pulmonary vascular damage biomarkers. E) The mRNA expression levels of key pulmonary vascular damage-related genes assessed and represented as above. Statistical differences in B), C), and E) were assessed by the Student’s t test. Significance is indicated as follows: ^**§**^p<0.05; ^**§§**^p<0.01 and ^**§§§**^p<0.001 infected+PBS treated versus uninfected mice; *p<0.05, **p<0.01 and ***p<0.001 infected+PBS treated versus infected+scFv76 treated mice.

### Structural bases for broad RBD recognition of SARS-CoV-2 variants by scFv76

To explore the recognition principles and to rationalize the broad cross-reactivity of scFv76 towards SARS-CoV-2 variants, we determined the 3D structure of the spike:scFv76 complex using single particle Cryo-EM. We used a SARS-CoV-2 Wuhan-Hu-1 6P-stabilized glycoprotein (Native Antigen) (*14*) incubated with scFv76 to assemble the complex. Our single particle cryo-EM analysis revealed a homogeneous population of the spike:scFv76 complex displaying two RBDs in the up conformation and one down, one scFv76 fragment being bound to the tip of each RBD (**Figure 5A)**.

**Figure 5.**
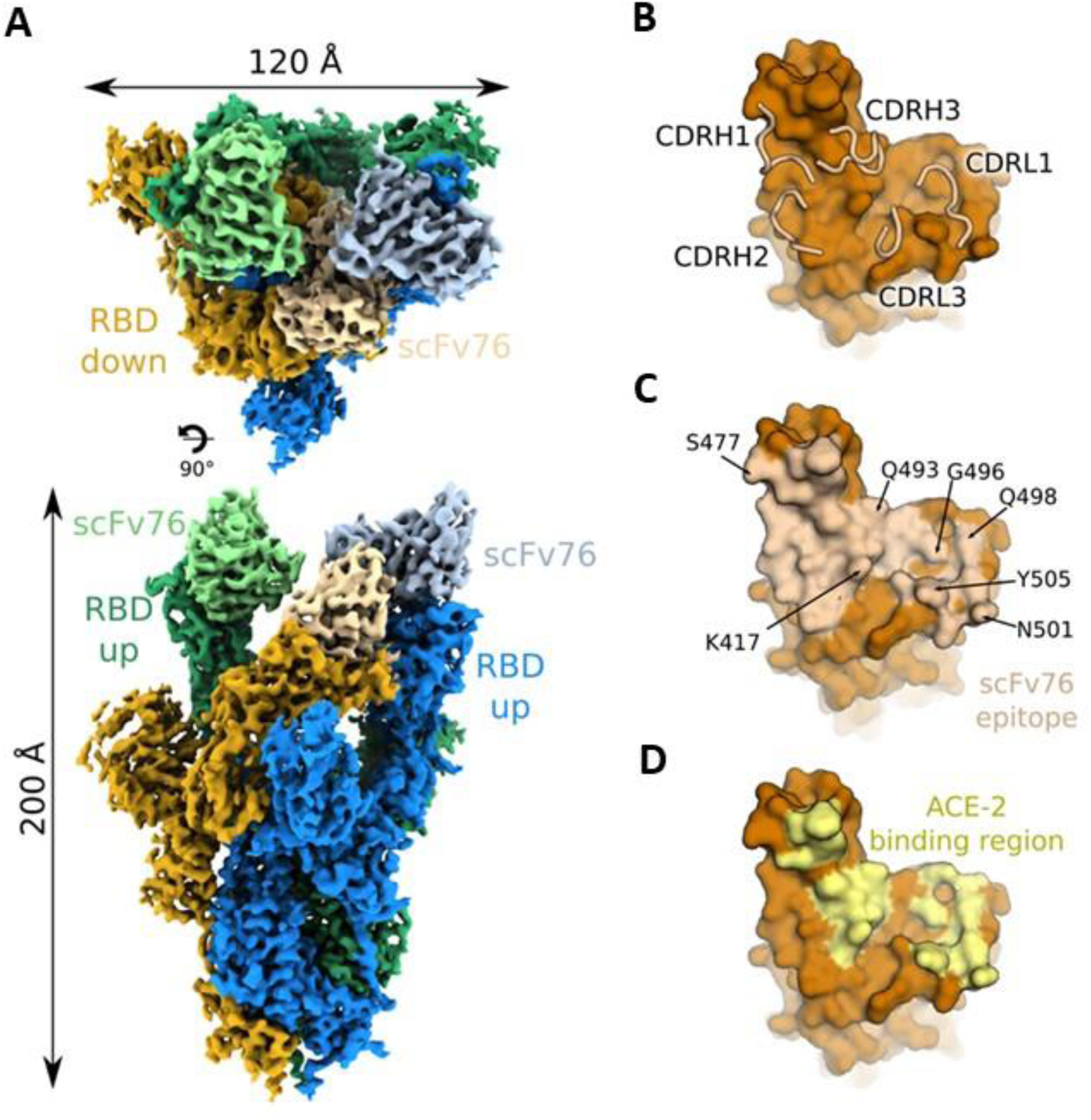
ScFv76 broad recognition of SARS-CoV-2 variants. A) Composite cryo-EM map of the SARS-CoV-2 spike protein with the locally refined RBD:scFv76 in two orientations. The RBD up or down conformations with their corresponding scFv76 fragments are labeled. The spike subunits are highlighted in green, blue, and yellow, respectively, and the scFv76 fragments bound to each RBD in light green, light blue and light yellow. B) ScFv76 CDR loops overlaid on surface representation of the RBD. C) RBD surface showing epitope residues as colored in B. D) RBD surface showing the ACE2 binding region colored in yellow.

The final 3D reconstruction displayed an overall resolution of 3.5 Å (**Figure S1A** and **Figure S2**); nevertheless, the epitope-paratope interface regions were less clearly resolved compared to the main spike component due to flexibility of the RBDs. To gain better insight into the recognition interface structure we applied a focused refinement procedure (*15*) on the RBD-down fragment region that brought the local resolution to 4.0 Å (**Figure 5A, Figure S1B** and **Figure S2**), and subsequently based our analysis on this structure. The scFv76:RBD refined structure showed that both the light and heavy chains of scFv76 contact the tip of the RBD, in both up and down conformations.

ScFv76 buries a surface area of ∼1082 Å^2^, based on an approximately equal contribution of the light and heavy chain components. The scFv76 paratope comprises residues of all three heavy chain complementarity determining regions (CDRs), and two from the light chain CDRs (**Table 3** and **Figure 5B**). Conversely, the recognition site lies on the RBD receptor-binding ridge and surrounding areas (**Figure 5C**), in full agreement with alanine scanning analysis previously reported (*12*), whereby RBD L455A, F456A, Y473A, N487A and Y489A mutations strongly reduced scFv76 binding. Of note, F456 is located within the deep groove created between CDRH1 and CDRH2, on one side, and CDRH3 and CDRL3 on the other. The scFv76:RBD interaction may also be stabilized by several hydrogen bonds (as evaluated on a 4.0 Å resolution structure). Among these, RBD D420 interacts with S56 (CDRH2), the carbonyl groups of RBD residues L455 and A475 interact with Y33 and T28 (both in CDRH1), respectively; the side chain of RBD Y421 falls close to P53 backbone carbonyl in CDRH2. Notably, the angle of approach of scFv76 to the RBD resembles that of ACE2; the scFv76 fragment contact region overlaps with the ACE2 binding interface, matching the location of 13 out of 17 ACE2-binding residues on RBD (**Figure 5D**). In this respect, the close resemblance of scFv76 and ACE2 binding modes to the RBD, and the ensuing competition for binding, explain on structural ground the potent scFv76 neutralizing activity. The scFv76:RBD pose resembles closely that observed for most antibodies from the VH3-53/VH3-66 germline. Nevertheless, not all such antibodies show neutralizing activity across SARS-CoV-2 variants, once more stressing the key role played by subtle and specific structural variations on the outcome of epitope-paratope interaction.

**Table 3.**
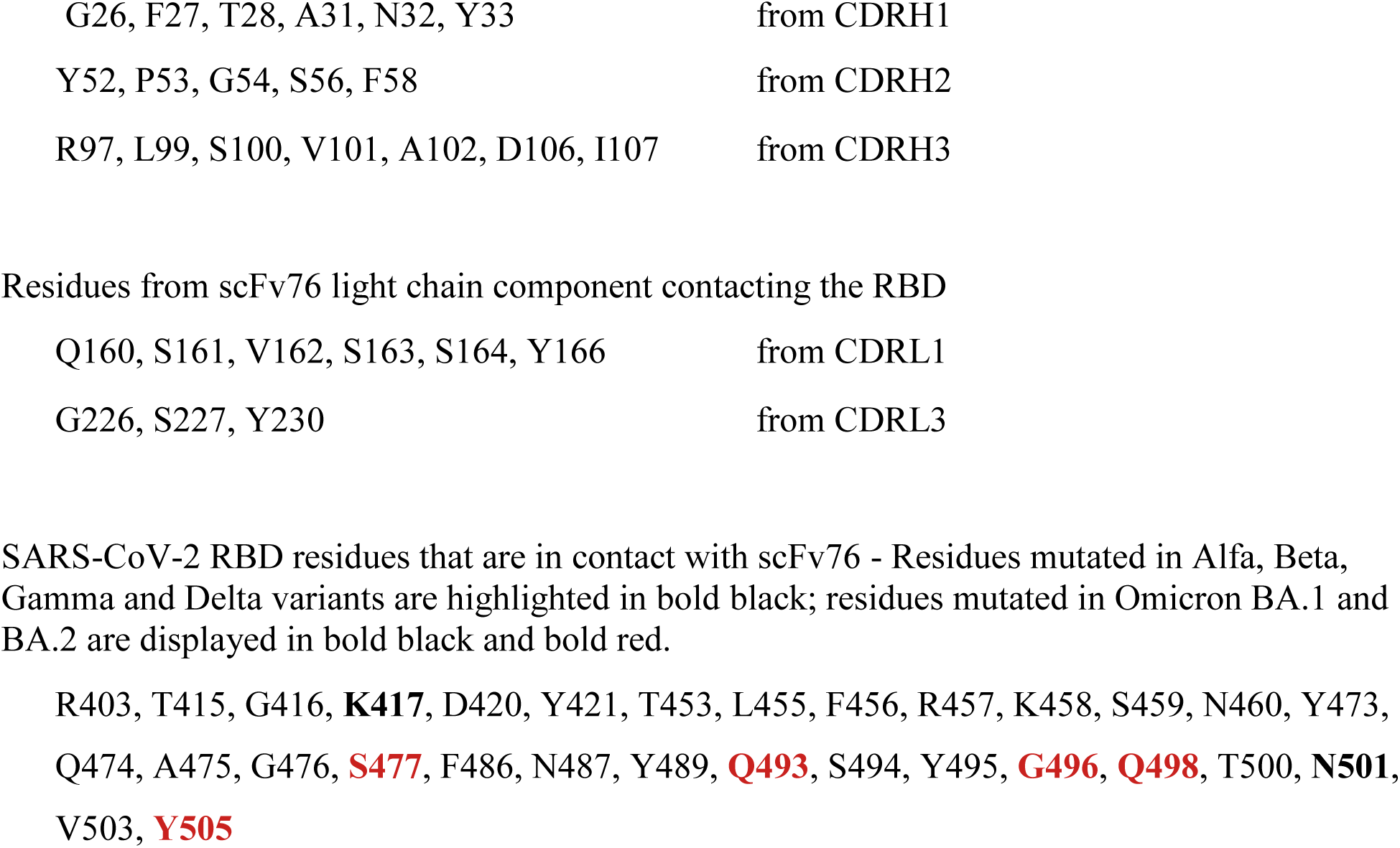
Residues from scFv76 heavy chain component contacting the RBD

|  |  |
| --- | --- |
| G26, F27, T28, A31, N32, Y33 | from CDRH1 |
| Y52, P53, G54, S56, F58 | from CDRH2 |
| R97, L99, S100, V101, A102, D106, I107 | from CDRH3 |
| Residues from scFv76 light chain component contacting the RBD |  |
| Q160, S161, V162, S163, S164, Y166 | from CDRL1 |
| G226, S227, Y230 | from CDRL3 |

SARS-CoV-2 RBD residues that are in contact with scFv76 - Residues mutated in Alfa, Beta, Gamma and Delta variants are highlighted in bold black; residues mutated in Omicron BA.1 and BA.2 are displayed in bold black and bold red.
R403, T415, G416, **K417**, D420, Y421, T453, L455, F456, R457, K458, S459, N460, Y473, Q474, A475, G476, **S477**, F486, N487, Y489, **Q493**, S494, Y495, **G496**, **Q498**, T500, **N501**, V503, **Y505**

A core of 28 epitope residues recognized by scFv76 is conserved in SARS-CoV-2 Alfa, Beta, Gamma and Delta variants carrying the K417N, E484K, and N501Y important mutations (**Table 3**). In our refined model E484 does not directly contact scFv76 consistently with previously shown reactivity of scFv76 with E484 mutated variants (*12*). We also predict a low susceptibility to mutations at K417 and N501 since they are not involved in any polar contact with scFv76. Moreover, both residues are in solvent exposed regions, allowing conformational flexibility: 417N would insert into a large groove created among CDRs, while 501Y would position the aromatic sidechain beyond the scFv76 CDRL3 (**Figure 5C** and **S3**). Both mutations are indeed known to have a limited impact on the scFv76 neutralizing power (*12*), in keeping with the broad SARS-CoV-2 recognition properties displayed by scFv76.

Notably, our modeling exercise suggests that the residues building the RBD epitope, recognized by scFv76, should drop to 23 in the Omicron BA.1 and BA.2 spike variants (**Table 3**). In fact, the five (Omicron-unique) sidechain substitutions, occurring within the RBD epitope region in these variants, are predicted to marginally affect scFv76 binding as confirmed by RBD/ACE2 competition and virus neutralization data, herein presented. Particularly, the S477N, Q493R, G496S and Q498R substitutions would place the mutated residues in solvent exposed regions (**Figure 5C** and **S3**). Residue Y505 is located between the CDRL1 and CDRL3 in the RBD:scFv76 complex, and mostly participates in hydrophobic contacts (**Figure 5C** and **S3**); the Omicron Y505H mutation may follow the same scheme. It is worth noting that such modeling considerations, supporting a substantial conservation of the RBD:scFv76 interface in all variants, are in agreement with the functional data reported in this paper that highlight the resilience of scFv76 to the main SARS-CoV-2 variants, including Omicron BA.1 and BA.2.

## DISCUSSION

In the search of easily deployable therapeutic measures against COVID-19, we recently described 76clAbs, a cluster of human single chain antibody fragments that, in principle, could bypass all limitations of traditional monoclonal antibodies. Indeed, the use of monoclonal antibodies for the therapy of COVID-19 is being challenged by several issues: 1) difficulties in the deployment of therapy, being monoclonal antibodies parenteral drugs to be administered in hospital environments; 2) the risk of antibody-dependent enhancement (ADE) that can be ignited by different routes involving the immunoglobulin Fc interaction with Fc receptor (*16*) or with the ACE2, recently found to possibly act as a secondary receptor (*17*), or with Fcγ-expressing cells including monocytes and macrophages that, by triggering the inflammatory cell death, needed to abort the production of infectious virus, cause systemic inflammation that contributes to the severity of COVID-19 pathogenesis (*18*); 3) evasion properties of SARS-CoV-2 variants, particularly recently emerged Omicron lineages for which most of approved and investigational antibodies lost their neutralization activity (*4-10*).

The single chain antibody format, because of its high stability, can be easily used for friendly self-administrable aerosol treatments. Furthermore, single-chain antibodies are, in principle, devoid of ADE risk because of lack of Fc sequence. 76clAbs, which were selected on the original SARS-CoV-2 Wuhan strain, were found to be resilient to Alfa, Beta, Gamma and Delta variant mutations (*12*). Importantly, we presently show that the scFv76 antibody of the cluster can also recognize and neutralize infectivity and fusogenic activity of Omicron BA.1 and BA.2 variants. Single particle cryo-EM results point to the peculiar property of this antibody to bind to both up and down conformation of the spike RBD, recognizing a well conserved epitope located at the ACE2 binding interface, and thus accounting for its neutralization properties. All mutations present in the RBD of the known SARS-CoV-2 variants are predicted to marginally impact scFv76 recognition, as confirmed by experimental results. In addition, we hypothesize that also the Omicron variants BA.4 and BA.5 might be still neutralized by scFv76. In fact, the spike mutated residues in these two variants, relative to Omicron BA.2, are 69-70del, L452R, F486V and wild type amino acid Q493 (*19*), being F486V the only mutation at the scFv76 binding interface. F486V replaces a bulky apolar side chain, closing a hydrophobic patch, with a smaller one, possibly affecting to a limited degree the stability of the complex.

Significant therapeutic efficacy of nebulized scFv76 is herein shown in a severe model of SARS-CoV-2 Delta pneumonia. The present data indicate that the aerosol treatment with scFv76 can efficiently control viral proliferation thus significantly reducing lung inflammation and damage. Altogether these results encourage further clinical development of the scFv76 antibody aerosol therapy as a new opportunity for the treatment of COVID-19, independently of the variant causing the disease.

## MATERIALS AND METHODS

### Spike/ACE2 binding competition

For competition experiments, Nunc MaxiSorp plates with 96 wells were coated with 100 μL/well of SARS-CoV-2 spike1 variant B.1.617.2 (Delta) protein (His Tag), SARS-CoV-2 spike S1+S2 trimer variant B.1.1.529 (Omicron) protein (ECD, His Tag), both from Sino Biological, and SARS-CoV-2 spike Trimer, variant BA.2 (Omicron) protein His Tag (MALS verified), from Acro Biosystems in PBS, at a final concentration of 0.5 μg/mL, overnight (ON) at +4 °C. Plates were blocked with 300 μL/well blocking solution for 2 h at room temperature (RT). After washing, dilutions of antibodies were added in a volume of 50 μL/well at double concentration, and after 30 min incubation at 37 °C, 1.0 μg/mL human ACE2 protein mouse Fc-tag (Sino Biological) was added and incubated for 1 h at 37 °C. Plates were washed 4 times with PBS/Tween and then incubated for 1 h at RT with 100 μL/well of an anti-mouse Fc conjugated to alkaline phosphatase (Sigma-Aldrich), diluted 1:1000 in blocking buffer. After 4 washings, 100 μL/well pNpp substrate were added and plates were incubated at RT in the dark. Absorbance was recorded at 405 nm using a SUNRISE Tecan spectrophotometer.

### Surface plasmon resonance

Kinetic constants were determined using SPR experiments with a Biacore T200 instrument (Cytiva). SARS-CoV-2 spike S1+S2 trimer variant B.1.1.529 (Omicron) protein and SARS-CoV-2 spike Trimer, variant BA.2 (Omicron) protein both 1.25 μg/mL in HBS-P+ buffer (Cytiva), were immobilized at 1000 RU level on the surface of a flow cell of a Series S sensor chip NTA (Cytiva) using a Ni^2+^-mediated capture followed by amine coupling procedure, while another flow cell surface was blank-immobilized by amine coupling with ethanolamine to be used as control surface. Then scFv76 was flowed at 30 μL/min on all flow cells at 0.47, 1.40, 4.19, 12.56, 37.67, and 113 nM concentrations in HBS-EP+ buffer (Cytiva) for a contact time of 480 s. After a dissociation time of 900 s, all flow cells surfaces were regenerated by flowing a solution of glycine-HCl 4 mM and SDS 0.1% w/w at 30 μL/min for 30 s. Double referenced sensorgrams were obtained by subtraction of blank-immobilized flow cell curves, as well as of zero concentration curves, from derivatized surface flow cell curves. Kinetic constants were obtained by BIAevaluation 3.2 software (Cytiva) fitting with a 1:1 binding model.

### Viral Neutralization in Calu-3 cells

To measure the SARS-CoV-2-neutralizing capability of scFv76, a live SARS-CoV-2 assay was performed by measuring the viral load in human lung adenocarcinoma Calu-3 cells, by real-time reverse transcription-quantitative PCR (RT-qPCR), 72 h after virus infection. The experiments were carried out at the François Hyafil Research Institute (Oncodesign; Villebon-sur-Yvette, France). Calu-3 cells were seeded in 96-well plates in complete cell culture medium (MEM + 1% pyruvate + 1% glutamine + 10% fetal bovine serum) and then infected, at a multiplicity of infection of 0.01, with SARS-CoV-2 Delta virus (provided by NIID Japan, strain hCoV-19/Japan/TY11-330-P1/2021; originally provided by the GISAID; accession ID: EPI_ISL_ 2158613). One hour after infection, the virus solution was discarded and replaced by a volume of growth medium containing scFv76 or not-neutralizing scFv5 antibody, in a concentration ranging from 214 to 2.6 nM, in triplicate. The plates were then transferred in a 37 °C incubator for 72 h. Finally, the cell culture supernatants were collected for viral RNA extraction (Macherey Nagel Viral RNA kit) and viral RNA copy number was quantified by RT-qPCR, targeting a region in the viral ORF1ab gene and using a QuantStudio™ 7 Real-Time PCR System (Applied Biosystems). Data were processed using GraphPad Prism software (V8.0) and the IC50 values calculated using a four-parameter logistic curve fitting approach.

### SARS-CoV-2 S-pseudovirus neutralization assay

Generation of SARS-CoV-2 S-pseudovirus and SARS-CoV-2 S-pseudovirus neutralization assay were performed as previously described (*12*). The vectors expressing the Omicron SARS-CoV-2-spike (S1+S2)-long (B.1.1.529) and SARS-CoV-2-spike (S1+S2)-long (B.1.1.529 sublineage BA.2) were obtained from GenScript and Sino Biological, respectively. Serial (1:3) dilutions (range: from 35.7 to 0.14 nM final concentration) of scFvs were tested in duplicate. Luciferase activity (Relative Luciferase Units or RLU) was detected at 72 h post-infection by using Bright-Glo Luciferase Assay System Kit (Promega) in a Microplate Luminometer (Wallac-Perkin Elmer).

### Microneutralization assay

Neutralizing antibody titers were tested using a live-virus assay as follows. ScFv samples were pre-diluted in inoculation medium (DMEM, 2% fetal calf serum, 1% Glutamine), followed by 9 serial dilutions in inoculation medium. Each serial dilution was then mixed 1:1 with 2,000 TCID50/mL SARS-CoV-2 variant virus (Delta variant: strain hCoV-19/USA/MD-HP05647/2021; Omicron variant: strain hCoV-19/USA/MD-HP20874/2021) and incubated for 1 h at + 37 °C ± 2 °C and 5% ± 0.5% of CO_2_. Thirty-five μL of each diluted sample/virus mix were then applied in octuplicate to Vero E6 cells seeded at a density of 10^4^ cells/well in a 96-well plate at day -1. After 1 h of incubation at +37 °C ± 2 °C and 5% ± 0.5% CO_2_, 65 μL of inoculation medium (DMEM, 2% fetal calf serum, 1% Glutamine) were added per well. Plates were incubated for six days at +37 °C ± 2 °C, 5% ± 0.5% CO_2_. After this incubation, the cells were inspected for cytopathic effects (CPEs) and the number of positive wells (that is exhibiting CPEs) recorded. Data were processed using GraphPad Prism software (V8.0) and the IC50 values calculated using a four-parameter logistic curve fitting approach.

### Cell-cell fusion assay

Human alveolar type II-like epithelial A549 cells and embryonic kidney 293T cells were obtained from ATCC (Manassas, VA). Cells were grown at 37 °C and 5% CO2, in RPMI-1640 (A549 cells) or DMEM (293T cells) medium (Euroclone) supplemented with 10% fetal calf serum (FCS), 2 mM glutamine, and antibiotics. Generation of A549 cells stably expressing the human ACE2 receptor (A549-hACE2 cells) has been described previously (*20*). The vectors expressing the Omicron SARS-CoV-2-spike (S1+S2)-long (B.1.1.529) and SARS-CoV-2-spike (S1+S2)-long (B.1.1.529 sublineage BA.2) were obtained from GenScript and Sino Biological, respectively. Transfections were performed using Lipofectamine 2000 (Invitrogen, Thermo Fisher Scientific), according to the manufacturer’s instructions. The donor-target cell fusion assay has been described previously (*12*). Transmission and fluorescence images were taken using a Zeiss Axio Observer inverted microscope and the extent of fusion was then quantified as described (*12*). Images shown in all figures are representative of at least five random fields (scale bars are indicated). Statistical analysis was performed using one-way ANOVA (Prism 6.0 software; GraphPad). All experiments were done in duplicate and repeated at least twice.

### *In vivo* pharmacological evaluation of nebulized scFv76 in a model of SARS-CoV-2 Delta pulmonary infection

The animal study was carried out at the San Raffaele Scientific Institute (Milan, Italy) and performed in accordance with the European Directive 2010/63/EU on the protection of animals used for scientific purposes, applied in Italy by the Legislative Decree 4 March 2014, n. 26. All experimental animal procedures were approved by the Institutional Animal Committee of San Raffaele Scientific Institute. Female transgenic K18-hACE2 mice, aged 8-10 weeks, were infected via the intranasal route with 1×10^5^ TCID50/mouse of SARS-Cov-2 variant Delta B.1.617.2 virus [hCoV-19/Italy/LOM-Milan-UNIMI9615/2021 (GISAID Accession ID: EPI_ISL_3073880)], obtained from the Laboratory of Microbiology and Virology of San Raffaele Scientific Institute. One hour and 8 h after infection, and twice/day for two additional days, infected mice (5/group) were treated by nose-only nebulization with 2.5 mL of scFv76 (3.0 mg/mL in PBS) or PBS by using an Aerogen Pro (Aerogen) mesh nebulizer and a nose-only inhalation chamber, suitable for delivering the nebulized antibodies contemporarily up to 8 mice, as previously described (*12*). Mice were monitored for appearance, behavior, and weight. At day 4 post infection they were euthanized by inhalation of 5% isoflurane followed by gentle cervical dislocation, and lungs and nasal turbinates were explanted and then fixed by 4% paraformaldehyde for IHC analyses or snap-frozen (in liquid nitrogen) and stored at -80 °C until further analyses.

### Tissue homogenization and viral titer determination

For viral titer determination in the lung, tissues homogenates were prepared by homogenizing perfused lungs using gentleMACS Octo Dissociator (Miltenyi) in M tubes containing 1 mL of DMEM. Samples were homogenized for three times with program m_Lung_01_02 (34 s, 164 rpm). The homogenates were centrifuged at 3,500 rpm for 5 min at 4 °C. The supernatant was collected and stored at -80 °C until use for viral isolation and viral load detection. Viral titer was calculated by 50% tissue culture infectious dose (TCID50). Briefly, Vero E6 cells were seeded at a density of 1.5 × 10^4^ cells per well in flat-bottom 96-well tissue culture plates. The following day, 2-fold dilutions of the homogenized tissue were applied to confluent cells and incubated for 1 h at 37 °C. Then, cells were washed with phosphate-buffered saline (PBS) and incubated for 72 h at 37 °C in DMEM 2% FBS. Cells were fixed with 4% paraformaldehyde for 20 min and stained with 0.05% (wt/vol) crystal violet in 20% methanol. The plate analysis was carried out by qualitative visual assessment of cytopathic effect. TCID50 was determined by Reed & Muench method.

### RT-qPCR for viral copies quantification and gene expression analysis

For viral copies quantification and gene expression analysis, tissues homogenates were prepared by homogenizing perfused lungs or nasal turbinates (NT) using gentleMACS dissociator (Miltenyi) with program RNA_02, in M tubes, in 1 mL or 500 μL Trizol (Invitrogen) for lungs or NT, respectively. The homogenates were centrifuged at 2,000 × g for 1 min at 4 °C, and the supernatant was then collected. RNA extraction was performed by combining phenol/guanidine-based lysis with silica membrane-based purification. Briefly, 100 μL of chloroform were added to 500 μL of homogenized sample; after centrifugation, the aqueous phase was added to 1 volume of 70% ethanol and loaded on ReliaPrep™ RNA Tissue Miniprep column (Promega, Cat #Z6111). Total RNA was isolated according to the manufacturer’s instructions. For viral copies quantification, quantitative polymerase chain reaction (qPCR) was performed using TaqMan Fast virus 1 Step PCR Master Mix (Applied Biosystems); a standard curve was drawn with 2019_nCOV_N Positive control (IDT), and the following primers and probe were used: 2019-nCoV_N1-Forward Primer (5’-GAC CCC AAA ATC AGC GAA AT-3’), 2019-nCoV_N1-Reverse Primer (5’-TCT GGT TAC TGC CAG TTG AAT CTG-3’), 2019-nCoV_N1-Probe (5’-FAM-ACC CCG CAT TAC GTT TGG TGG ACC-BHQ1-3’) (Centers for Disease Control and Prevention (CDC) Atlanta, GA 30333). All experiments were performed in duplicate.

For gene expression analysis of inflammation and endothelial-related genes, total RNA was retrotranscribed using the SuperScript IV VILO Mastermix (Invitrogen; Thermo Fisher Scientific), according to the manufacturer’s instructions. Real time qPCR analysis was performed using TaqMan Fast Advanced Master Mix and specific TaqMan Gene Expression Assays (listed in **Table S1**), both from Applied Biosystems (Thermo Fisher Scientific). The 7900HT Sequence Detection System instrument and software (Applied Biosystems) were used to quantify the mRNA levels of the target genes, according to a six-point serial standard curve generated for each gene. Results were ultimately expressed, after normalization to the housekeeping gene Rlp32, as relative expression (fold change) as compared to uninfected animals.

### Histopathological analysis

PBS-perfused lungs were fixed in Zn-formalin for 24 h and then stored in 70% ethanol until trimming for paraffin wax embedding and the following histological examination. Consecutive sections (20 μm) were prepared and stained by the classical hematoxylin & eosin (H&E) method, and then microscopic observation was performed using a Nikon Eclipse 80i microscope equipped with a DXM1200F Microscope Camera. Pathological features in lung sections were scored as follows: inflammation-related parameters, including congestion of the alveolar septa, lymphomonocyte interstitial (alveoli) infiltrate, alveolar hemorrhage, interstitial edema and platelet microthrombi, were separately evaluated by two independent pathologists and the extent of these findings was arbitrarily scored, using a two-tiered system, as 0 (negative), 1 (moderate) and 2 (severe). All the scores of each animal were ultimately summed up and a global score was calculated for each group and expressed as the average ± SE. The percentage of pulmonary area affected by lesions in each section was also measured and results for each group were reported as average percentage ± SE.

### Electron microscopy sample preparation

A sample of SARS-CoV-2 Wuhan-Hu-1 6P-stabilized glycoprotein (Native Antigen) was incubated with scFv76 at a final concentration of 0.65 mg/mL and 0.22 mg/mL, respectively, for 1 h at RT. A 4 μL droplet of the sample was applied onto a R1.2/1.3 300-mesh copper holey carbon grid (Quantifoil), previously glow discharged for 30 s at 30 mA using a GloQube system (Quorum Technologies). The sample was incubated on grid for 60 s at 4 ºC and 100% relative humidity, blotted and plunge-frozen in liquid ethane using a Vitrobot Mk IV (Thermo Fisher Scientific).

### EM Data collection and image processing

Cryo-EM data were acquired on a Talos Arctica (Thermo Fisher Scientific) transmission electron microscope operated at 200 kV. The data were acquired using EPU-2.8 automated data collection software (Thermo Fisher Scientific). Images were collected at nominal magnification of 120,000×, corresponding to a pixel size of 0.889 Å/pixel at the specimen level, with applied defocus values between -0.8 and -2.2 μm. A total of 4211 movies were acquired using the Falcon 3 direct electron detector (Thermo Fisher Scientific) operating in electron counting mode, with a total accumulated dose of 40 e^-^/A^2^ distributed over 40 movie frames.

Movies were preprocessed with WARP 1.0.9 (*21*). A 5×5×40 model was used for motion correction using a 35-7 Å resolution range weighted with a -500 Å^2^ B factor. Contrast transfer function (CTF) was estimated using the 40-3.5 Å resolution range and a 5×5 patch model. Particle picking was performed using the deep convolutional neural network BoxNet2Mask_20180918, resulting in 490,614 particles that were extracted in 400-pixel boxes and imported into CRYOSPARC-3.3.1 *(15)* for further processing.

2D classification was used to select 366,967 particles that were 3D aligned using the SARS-CoV-2 spike glycoprotein model (EMDB-21452) low pass filtered at 30 Å as reference. Particles were subjected to 3D classification to select the final set of 87,623 particles that yielded an overall 3.5 Å resolution reconstruction based on the gold-standard criterion of 0.143 Fourier shell correlation (FSC) value. The initial reconstruction displayed two spike RBDs in up conformation and one down, all three showing one bound scFv76 fragment. Particles were subtracted with a mask comprising the entire spike molecule without the RBD in the down conformation and its corresponding scFv76 fragment, and then locally refined to 4.0 Å resolution according to FSC of 0.143.

### Model building, refinement and validation and structural analysis

The scFv76 structure was modelled with Antibody Structure Prediction module using the Schrö-dinger Maestro Bioluminate Suite 4.5.137, release 2021-4 (*22*). The modelling was performed with antigen-binding fragment (Fv) as antibody format. We used “EVQLLQSAGGLVQPGGSLRLS- CAASGFTVSANYMSWVRQAPGKGLEWVSVI- YPGGSTFYADSVKGRFTISRDNSKNTLYLQMNSLRVEDTAVYYCARDLSVAGAFDIWGQG TLVTVSSGG” as target sequence for the heavy chain (HC) and “IVLTQSPGTLSLSPGER- ATLSCRASQSVSSSYLAWYQQKPGQAPRLLIYGASSRATGIP- DRFSGSGSGTDFTLTISRLEPEDFAVYYCQQYGSSPYTFGQGTKLEIKRAAAGDYK” for the light chain (LC). The tool identifies the best matching framework templates for the queried sequences and builds the CDR loops based on the cluster analysis performed on the default antibody loop database, sieved using the selected framework template. To select a suitable structural reference framework, we filtered the possible candidates according to the scFv76 HC and LC germlines, IGHV3-66 and IGKV3-20, respectively (*12*). We analyzed 130 CoV-AbDab structures of antibodies (Abs) bound to RBDs *(23)* and found that 39 out of 42 entries with IGHV3-53/IGHV3-66 HCs, usually coupled with IGKV1-9 (16 Abs) or IGKV3-20 (10 Abs) LCs, share a common binding mode to the RBD. The three outliers are characterized by longer CDR-H3 (17-25 residues compared to 8-15 residues) and they are bound to IGLV2-14/IGLV2-23 LCs. Based on these findings and considering the good resolution (2.03Å) and the average of heavy and light chain similarity scores (0.99 out of 1.00), we selected PDB entry 7N3I as the reference framework, whose HC and LC are a combination of IGHV3-53 and IGKV3-20. To model the scFv76 CDR loops, as well as their interaction with the antigen, the CDRs were grafted into a homology modelled structure built based on the reference framework template that also included the N-terminal domain of the betacoronavirus-like trimeric spike glycoprotein S1 (pdb 7N3I). The generated scFv76:RBD model was subsequently superimposed on the three RBDs of a SARS-CoV-2 HexaPro S cryo-EM structure, with two RBDs in up and one in down conformation (PDB 7N0H). Finally, the three cryo-EM RBDs in complex with the modelled scFv76 were refined with Schrödinger Protein Preparation Wizard (*24*) to remove clashes and optimize sidechains.

The model generated was split into two parts: the first comprised the whole spike protein without the RBDs; the second consisted of the RBD in the down conformation in complex with scFv76. Both models were independently refined with COOT (*25*) and PHENIX (*26*) using the full reconstruction and the local refined map, at 3.4 Å and 4.0 Å resolution, respectively. Subsequently, the local refined RBD:scFv76 complex was rigid body fitted in the full spike:scFv76 reconstruction. All data collection, image processing and final model statistics are summarized in **Table S2**. The images were prepared using ChimeraX (*27*) and Pymol (http://www.pymol.org/pymol).

## Statistical analysis

Statistical analyses were performed using Prism software (version 6.0 or 8.0 GraphPad Software). Data were presented as average ± standard error (SE) or ± standard deviation (SD). Statistical significance was analyzed with unpaired two-tailed Student’s t test, Mann-Whitney U-test or oneway analysis of variance (ANOVA). P values below 0.05 were considered statistically significant.

## Acknowledgements

We thank Prof Alessandro Rambaldi (Bergamo Hospital and Milan University) for suggestions and encouragement, the San Raffaele team for the effort and dedication devoted to the aerosol treatment of Delta-infected mice, Bruno Bruni Ercole and Evelyn Vaccaro (Alfasigma) for excellent technical assistance.

## Funding

This work was funded by Alfasigma SpA.

## Author contributions

Conceptualization: R.D.S., F.M.M. and O.M. Investigation: F.M.M., O.M., D.S., A.M.A., G.B., C.C., A.R.^1^, C.A., E.M., L.L., C.V., A.R.^5^, A.R.^6^, F.B., A.C-S., C.P. and C.C.^7^. Writing: F.M.M., R.D.S and M.B. Supervision: R.D.S., M.B., E.M.P, L.G.S. and M.G.S.

## Competing interests

O.M., E.M.P. and R.D.S. are employees of Alfasigma SpA and are named as inventors in a patent application on the name of the same Company. Other authors declare that they have no competing interests.

## Data and materials availability

Antibodies described in the paper can be provided upon material transfer agreement (MTA) subscription. The full spike:scFv76 and the RBD:scFv76 cryo-EM volumes and the structure coordinates have been deposited in the Electron Microscopy Data Bank and the Protein Data Bank under accession codes EMD-14628 and EMD-14629, and 7ZCE and 7ZCF, respectively. Cryo-EM movies were deposited in the Electron Microscopy Public Image Archive under the accession code EMPIAR-10990.

