## Supplementar tables and figures for "Spike mutation resilient scFv76 antibody counteracts SARS-CoV-2 lung damage upon aerosol delivery"

**Table S1.** TaqMan gene assays (from Thermo Fisher Scientific) used for real time qPCR analyses.

| Assay ID | Gene Symbol |
|---|---|
| Mm 00657574_s1 | **ANG2** |
| Mm00441242_m1 | **Ccl2** |
| Mm01268754_m1 | **Ccl20** |
| Mm00486938_m1 | **Cdh5** |
| Mm03294838_g1 | **COX-2** |
| Mm01290062_m1 | **Csf2** |
| Mm04207460_m1 | **Cxcl1** |
| Mm00445235_m1 | **Cxcl10** |
| Mm00438258_m1 | **Cxcr2** |
| Mm00516023_m1 | **ICAM-1** |
| Mm00515153_m1 | **Ifit1** |
| Mm03030145_gH | **IFNA1** |
| Mm00439552_s1 | **IFNB1** |
| Mm01168134_m1 | **IFNG** |
| Mm00439620_m1 | **IL1A** |
| Mm00434228_m1 | **IL1B** |
| Mm00445259_m1 | **IL4** |
| Mm00446190_m1 | **IL6** |
| Mm01288386_m1 | **IL10** |
| Mm00517640_m1 | **IL21** |
| Mm01705338_s1 | **lSG15** |
| Mm00487796_m1 | **MX1** |
| Mm00840904_m1 | **NLRP3** |
| Mm02528467_g1 | **Rlp32** |
| Mm00441278_m1 | **SELE** |
| Mm00443258_m1 | **TNF-a** |
| Mm01320970_m1 | **VCAM1** |

**Table S2.** Cryo-EM data collection, image processing and model refinement statistics.

| | | |
|---|---|---|
| **Data collection and image processing** | | |
| Structure | Spike:scFv76 full complex | RBD:scFv76 complex (Focused refinement) |
| Microscope | Thermo Fisher Scientific TALOS Arctica | |
| Voltage (kV) | 200 | |
| Camera | Falcon 3EC | |
| Magnification | × 120,000 | |
| Total electron dose ($e^-$/Å$^2$) | 40.0 | |
| Defocus range (μm) | -0.8 and -2.2 μm | |
| Pixel size (Å) | 0.889 | |
| Micrographs (no.) | 4,211 | |
| Symmetry imposed | C1 | C1 |
| Initial particle images (no.) | 490,614 | 87,623 |
| Final particle images (no.) | 87,623 | 87,623 |
| Resolution (Å) | 3.5 | 4.0 |
| (FSC threshold) | (0.143) | (0.143) |
| Sharpening B-factor (Å$^2$) | -136.6 | -157.9 |
| EMDB code | EMD-14628 | EMD-14629 |
| **Model refinement** | | |
| Protein residues | 3706 | 426 |
| N-acetyl-D-glucosamine molecules | 51 | 1 |
| r.m.s. deviations | | |
| Bond lengths (Å) | 0.004 | 0.002 |
| Bond angles (°) | 0.663 | 0.643 |
| Ramachandran plot | | |
| Favored (%) | 91.42 | 85.24 |
| Allowed (%) | 8.58 | 14.76 |
| Disallowed (%) | 0.00 | 0.00 |
| Validation | | |
| Molprobity score | 1.96 | 2.18 |
| Clashscore | 8.35 | 10.15 |
| Poor rotamers (%) | 0.03 | 0.56 |
| Map-model correlation | 0.84 | 0.72 |
| PDB code | 7ZCE | 7ZCF |

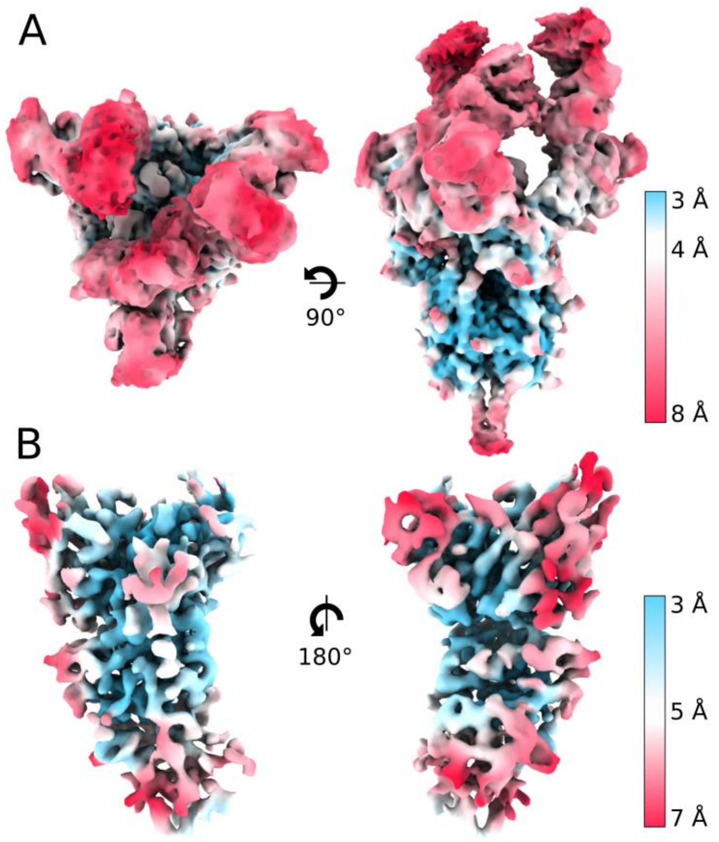


**Figure S1. Local resolution cryo-EM map.** A) spike:scFv76 full complex top and sides views. B) Locally refined RBD:scFv76 in the closed conformation, front and back views. Maps are colored according to the estimated local resolution.

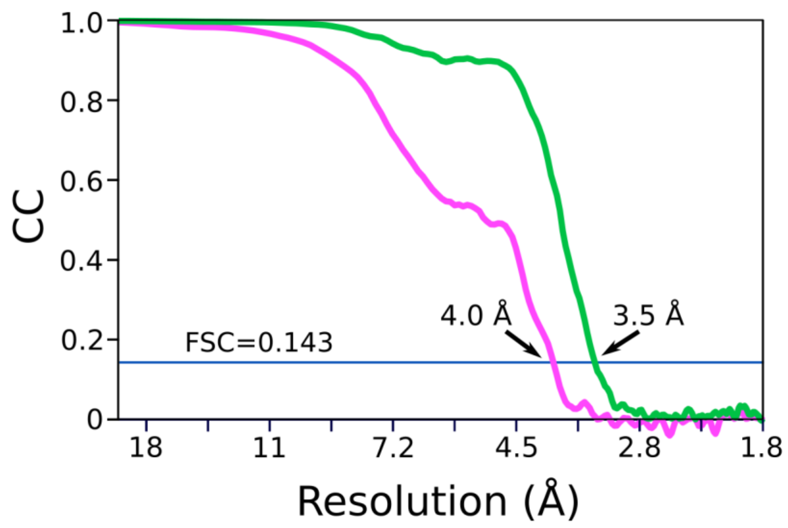


**Figure S2. Cryo-EM FSC curves of spike:scFv76.** The FSC curves for the spike:scFv76 full complex and the locally refined RBD:scFv76 in the closed conformation are shown in green and pink, respectively. The 0.143 threshold and the resolution cut-off for each curve are indicated.

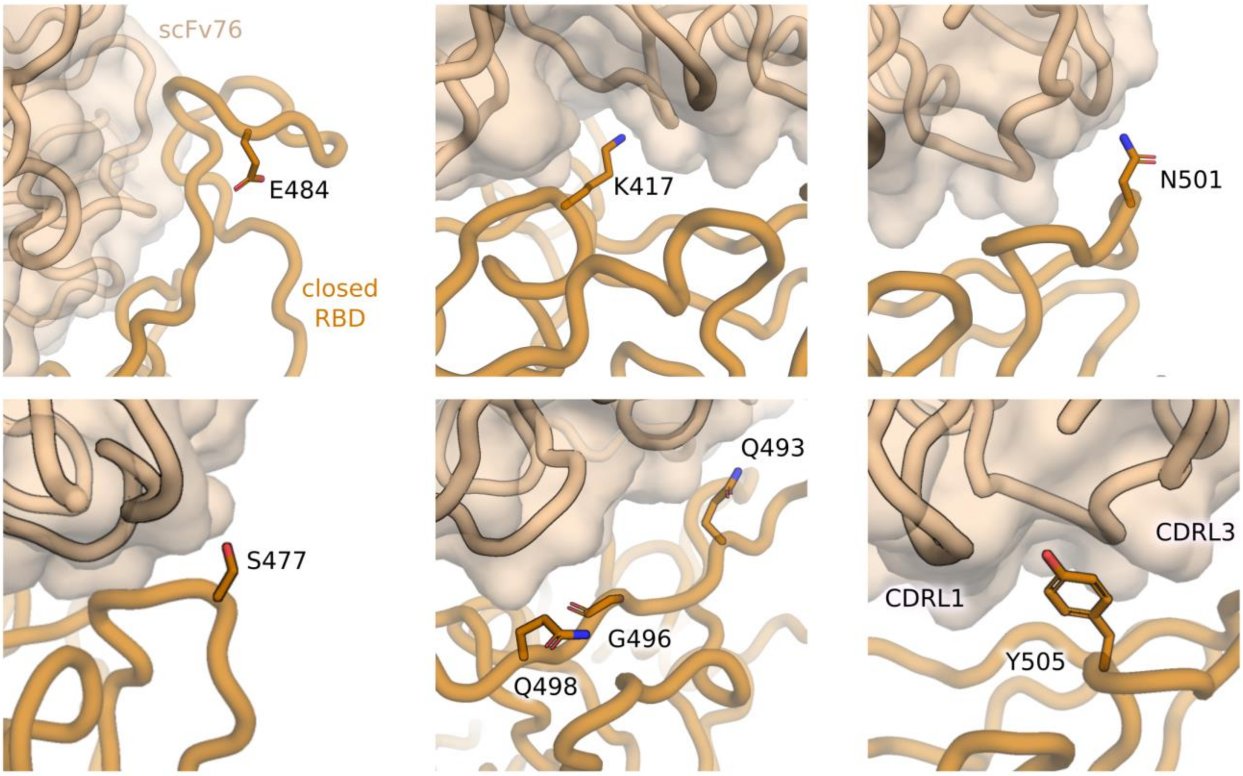


**Figure S3. SARS-CoV-2 variants mutations at the RBD:scFv76 interface.** Cartoon representation of scFv76 (transparent surface and worm) bound to the closed RBD (worm model); highlighted are key mutated residues (shown as stick models) in SARS-CoV-2 Omicron BA.1 and BA.2 variants.
